# Age-Related Changes in the Neural Dynamics of Bottom-Up and Top-Down Processing During Visual Object Recognition: An Electrophysiological Investigation

**DOI:** 10.1101/608240

**Authors:** Leslie Y. Lai, Romy Frömer, Elena K. Festa, William C. Heindel

## Abstract

When recognizing objects in our environments, we rely on both what we see and what we know. While elderly adults have been found to display increased sensitivity to top-down influences of contextual information during object recognition, the locus of this increased sensitivity remains unresolved. To address this issue, we examined the effects of aging on the neural dynamics of bottom-up and top-down visual processing during rapid object recognition. Specific EEG ERP components indexing bottom-up and top-down processes along the visual processing stream were assessed while systematically manipulating the degree of object ambiguity and scene context congruity. An increase in early attentional feedback mechanisms (as indexed by N1) as well as a functional reallocation of executive attentional resources (as indexed by P200) prior to object identification were observed in elderly adults, while post-perceptual semantic integration (as indexed by N400) remained intact. These findings suggest that compromised bottom-up perceptual processing of visual input in healthy aging leads to an increased involvement of top-down processes to resolve greater perceptual ambiguity during object recognition.

---

Real world objects are rarely seen in the absence of any contextual information. Rather, our day-to-day visual experience includes exposure to multiple types of normative or expected contextual relationships between object and scene, including spatial-physical relations (e.g., a computer monitor is normally on a desk rather than beneath it), semantic associations (e.g., a piano is in the living room rather than the kitchen), relative spatial orientations (e.g., chairs are normally oriented toward the table; cars are oriented towards the driving direction of a street), and scene probability (e.g., a pillow fight happening on the street is less probable than one happening in a bedroom). There is now substantial evidence that contextual information that is consistent with our knowledge, expectation, or visual experience facilitates object recognition (see Oliva & Torralba, 2007; Bar, 2004 for reviews), while contextual information that violates expected relationships impairs recognition (Biederman, Glass, & Stacy, 1973; Joubert, Rousselet, Fize, & Fabre-Thorpe, 2007, 2008; Fize, Cauchoix, & Fabre-Thorpe, 2011). The important role of context in guiding the perceptual hypotheses elicited by bottom-up sensory signals during object recognition (Trapp & Bar, 2015) suggests that object recognition involves close dynamic interactions between bottom-up and top-down processes. Indeed, top-down contextual modulation has been found to be available in the earliest stages of the recognition processes and prior to identification (Auckland et al., 2007, Davenport & Potter, 2004; Davenport, 2007). For example, it has been shown that observers can, within a glimpse, report more detail, perform better classification and show greater sensitivity to scene images that are consistent with prior visual experience (Greene, Botros, & Beck, 2015). An analysis of eye movements within a rapid visual categorization paradigm further suggests that contextual interference with object processing can begin as early as 160 ms following stimulus presentation (Crouzet, Joubert, Thorpe, & Fabre-Thorpe, 2009).

Event-related potential (ERP) studies have begun to identify some of the neural correlates of top-down processes mediating object recognition. Scene congruity effects have been consistently observed with the N300/N400 complex, with incongruent object-scene semantic relations associated with a greater negative deflection than congruent relations (Ganis & Kutas, 2003; Mudrik, Lamy, Deouell, 2010; Sun, Simon-Dack, Gordon, & Teder, 2011; Vo & Wolfe, 2013). The N300/N400 congruity effect can be elicited even when the mismatch between object and scene occurs prior to fixation on the object, suggesting that semantic information is available in extrafoveal vision and can serve as a source of guidance for fixation selection (Coco, Nuthmann, & Dimigen, 2018). Several studies have also reported knowledge-based contextual effects on earlier ERP components beginning around 100 ms that are associated with visual processing (Abdel Rahman & Sommer, 2008; Dambacher, Rolfs, Gollner, Kliegl, & Jacobs, 2009; Mudrik, Shalgi, Lamy, & Deouell, 2014; Tanaka & Curran, 2001). Context congruity effects within an object categorization task have been found to occur around 170 ms followed by the modulation of the N300/N400 complex (Guillaume, Tinard, Baier, & Dufau, 2018), suggesting that bottom-up perceptual processes interact continuously with top-down influences throughout the visual processing stream prior to object identification (Rao & Ballard, 1999; Bar, 2004; Trapp & Bar, 2015).

Changes in visual processing associated with healthy aging may alter the dynamics of bottom-up and top-down processes during object recognition. Previously identified age-related changes in visual processing include early sensory and perceptual loss (see Owsley, 2011; Habak & Faubert, 2000 for reviews), increased internal noise and decreased signal-to-noise ratio within primary visual cortex (Arena, Hutchinson, Shimozaki, & Long, 2013; Schmolesky, 2003), and neural dedifferentiation (i.e. reduced distinctiveness of neural representation) within object selective areas in the ventral visual stream (Park et al., 2004, 2011; Chee et al., 2006; Payer et al., 2006; Voss et al., 2008). Age-related alterations in top-down processes have also been identified, including impairments in the top-down inhibition of irrelevant information within visual areas (Gazzaley et al., 2008; Zanto, Toy, & Gazzaley, 2010). Despite these documented changes, however, the effects of aging on the interactive mechanisms mediating object recognition are still not well understood. Here, we postulate that age-related generalized slowing of information processing (Salthouse, 1996) along with deficits in sensory and perceptual processing (Owsley, 2011) diminish the efficiency of bottom-up processing, which in turn biases the aging visual system toward additional top-down processing to solve object recognition.

A few behavioral and electrophysiological studies have begun to identify effects of aging on bottom-up and top-down processes mediating object recognition that are consistent with this view. During a rapid object categorization task, older adults demonstrated a selective impairment for objects that were presented either peripherally or with low contrast (Lenoble, Bordaberry, Rougier, Boucart, & Delord, 2013), suggesting an age-related decline in the efficiency of feedforward processing within the ventral visual stream that is critical for rapid categorization (Serre, Oliva, & Poggio, 2007). Older adults also show greater effects of object-scene congruity than young adults during rapid object categorization (Remy et al., 2013), suggesting an increase in top-down engagement during visual recognition with age. EEG studies investigating object processing in older adults, typically employing stimuli consisting of simple shapes or faces (e.g. Stormer, Li, & Heekeren, 2013; Eimer, 2000; Rossion et al., 2000), have also found that early attentional mechanisms associated with sensory and perceptual processing may be altered in healthy aging. Both the amplitude and the latency of early ERP components sensitive to attentional mechanisms, such as the N1 and P200 (Potts, Patel, & Azzam, 2004; Doran & Hoffman, 2010; Drew, McCollough, Horowitz, & Vogel, 2009), have been found to be impacted by aging (Störmer et al., 2013; Gazzaley et al., 2008; Zanto et al., 2010; Bourisly & Shuaib, 2018).

Taken together, these findings suggest that compromised bottom-up perceptual processing of visual input in healthy aging leads to an increased involvement of top-down processes to resolve greater perceptual ambiguity during object recognition. In young adults, context-related top-down processing has been found to facilitate recognition under conditions of ambiguity by prioritizing relevant interpretations elicited by bottom-up signals (Trapp & Bar, 2015; Bar, 2004; Bar et al., 2006; Dyck & Brodeur, 2015). Consistent with the presence of increased perceptual ambiguity, older adults have been found to display increased electrophysiological activity associated with orbitofrontal cortex (OFC) around 100 ms post stimulus onset relative to young adults during an odd-ball detection task (Kaufman, Keith, & Perlstein, 2006), and dynamic causal modeling suggests that older adults’ electrophysiological signals recorded during a visual object naming task are driven by increased prefrontal inputs relative to young adults (Gilbert & Moran, 2016). Thus, to the degree that rapid top-down feedback signals from OFC to V1 reflect the generation of top-down predictions that subsequently bias bottom-up processing during recognition (Bar, 2004; Fenske, Aminoff, Fronau, & Bar, 2006; Kveraga, Boshyan, & Bar, 2007), these findings support the presence of an age-related bias toward top-down processing during object recognition.

The aim of the present study was to better understand the effects of aging on the electrophysiological signals related to the dynamics between bottom-up and top-down visual processing during a rapid object recognition task. We manipulated the strength of bottom-up and top-down signals of the input images by systematically varying the degree of object ambiguity and context congruity, respectively, and then examined the impact of these manipulations on specific ERP components indexing bottom-up and top-down processes mediating object recognition at different points along the visual processing stream. In particular, we examined the effects of object ambiguity and scene congruity on the N1 as an index of early sensory and attentional processing during visual discrimination, the P200 as an index of pre-perceptual top-down selective attention processes, and the N400 as an index of post-perceptual semantic integration processes. Based on previous studies in young adults, we expected to observe scene congruity effects in the first 200 ms time window in posterior regions (Guillaume et al., 2016) as well as in the 300-500 ms time window in fronto-central regions (Ganis & Kutas, 2003; Mudrik et al., 2010), and to see object perceptual ambiguity effects on ERP components during the first 200 ms time window. Finally, in line with the evidence that age-related increases in prefrontal engagement is present starting at 100 ms post stimulus onset, we expected to see age-related alterations in the degree of N1 and P200 modulation by bottom-up (i.e. object ambiguity) manipulations, reflecting age-related changes in early interactions between attentional and perceptual processing.

## METHODS

### Participants

Thirty healthy young participants (18-34 years of age; 11 males) and thirty healthy elderly participants without cognitive impairment (60-84 years of age; 11 males) participated in this study. Young participants were recruited from the student population (undergraduate and graduate) of Brown University, and elderly participants were recruited from the Rhode Island community. All participants provided written consent prior to participation in the study, and received either course credit or monetary compensation for their time. The protocol was approved by the Institutional Review Board at Brown University in accordance with the Helsinki Declaration.

All young participants reported normal or corrected-to-normal vision. Five elderly participants reported having cataracts and/or glaucoma, with only one having not had corrective surgery. All elderly participants had normal or corrected-to-normal visual acuity as assessed by the Low-Contrast Tumbling “E” Translucent Eye Chart. Five elderly participants scored within the moderate range of the Mars Letter Contrast Sensitivity Test while the rest of the elderly participants scored within the normal range. No elderly participant reported a history of cognitive impairment, and they all scored within the cognitively-normal range of the Mini-Mental State Examination (MMSE) and the Repeatable Battery for the Assessment of Neuropsychological Status (RBANS) (Randolph, Tierney, Mohr, & Chase, 1998). Demographic data for all participants, and the vision assessment, MMSE and RBANS results for the elderly participants are presented in Table 1.

**Table 1.** Mean values for demographic information, visual assessments, and neuropsychological tests.

|  | **Young Group**<br>**(n = 30)** | **Elderly Group**<br>**(n = 30)** |
| --- | --- | --- |
| Age, years (SD) [$t = -2.53, p = .01$] | 22.30 (3.91) | 70.03 (6.47) |
| Sex (male/female) [$\chi^2 = 0, p = 1$] | 11/19 | 11/19 |
| Education, years (SD) [$t = -34.56, p < .01$] | 15.10 (2.02) | 16.37 (1.85) |
| Tumbling 'E' Visual Acuity Test (SD) | - | 21.65 (5.26) |
| Mars Contrast Sensitivity Test (SD) | - | 1.66 (0.17) |
| MMSE (SD) | - | 29.30 (0.79) |
| RBANS Total Score (SD) | - | 104.07 (13.04) |

Note: The mean Tumbling 'E' test score is equivalent to an average visual acuity between 20/20 and 20/25 for elderly adults at 100% contrast level. The mean binocular contrast sensitivity score is within the normal range (1.52 - 1.76) for adults over the age of 60. MMSE = Mini-Mental State Examination; RBANS = Repeatable Battery for the Assessment of Neuropsychological Status

### Apparatus & Stimulus

The stimuli consisted of a set of 96 real-world photographs selected from a database used in a previous study (Greene, Botros, Beck & Fei-Fei, 2015), with 48 photographs depicting high-probability real-world scenes (e.g., cyclist on the road) and the other 48 depicting low-probability real-world scenes (e.g. under water conference). Improbable and probable photographs were matched as closely as possible in style, content, and structure, as well as low-level visual features (see Greene et al., 2015 for details). To manipulate object ambiguity, we created a blurred version of each of the 96 photographs from the set by applying a Gaussian filter of 1.5 pixel radius to the target object defined in each scene using Adobe Photoshop. Examples of the four stimulus conditions are shown in Figure 1a. The stimuli measuring 500 × 500 pixels were displayed at the center of the screen using Matlab R2017b (The MathWorks Inc., Natick, MA) and Psychophysics Toolbox extensions (ver. 3.0.13; Brainard, 1997; Pelli, 1997; Keiner et al., 2007) on a 24.1 inch ASUS LCD monitor (resolution 1920 by 1200) with a 60 Hz refresh rate. Participants viewed each stimulus image at a distance of approximately 77 cm from the monitor, subtending 9.82° by 9.82° visual angle.

**Figure 1.**
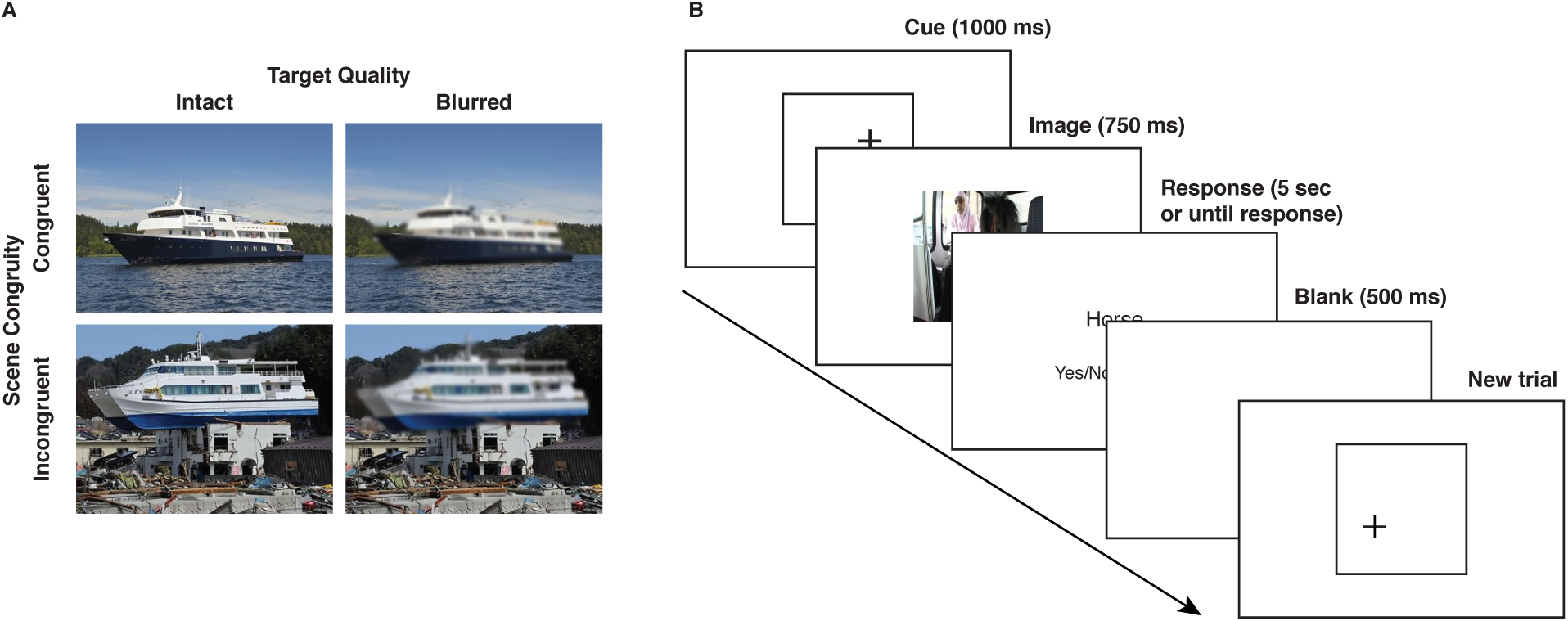
Examples of the four stimulus conditions (a). Main task trial illustration (b).

### Task & Procedure

The testing session included the main task and an eye movement recording task, both with EEG recordings. In the main task (Figure 1b), each trial began with a fixation cross presented at a variable location inside a bounding box (circumscribing the location of the to-be-presented visual scene) measuring 500 x 500 pixels at the center of the screen. Participants were informed that the fixation cross indicated the target location within the upcoming scene, and were instructed to fixate on the fixation cross while it remained on the screen. After a delay of 1 second, the bounding box along with the fixation cross was replaced by a trial image presented for 750 ms, followed by a response screen displaying a target word. Participants indicated whether the target word matched the target object they saw in the trial image by pressing one of the three response keys (i.e. ‘Yes’, ‘No’ or ‘Unsure’) as quickly and accurately as possible. Immediately following a response or after 5 seconds had elapsed following the stimulus onset, a blank screen of 500 ms was presented prior to the onset of the next trial.

Participants first completed a block of 20 practice trials without receiving performance feedback. If the practice block *Accuracy* was lower than 80%, the experimenter re-explained the task instructions and restarted the practice trials. After the practice block(s), each participant completed a single block of 96 experimental trials presented in randomized order. In half of the experimental trials (48 trials), target objects were presented intact whereas the other half presented blurred target objects. Half of the intact and blurred targets were presented within probable scenes whereas the other half were presented within improbable scenes. Target quality (i.e. intact or blurred) and scene congruity (i.e. probable or improbable) were randomized across trials and across participants. At the end of the testing session, participants completed an eye movement recording task that measured prototypical eye movements and blinks to serve subsequent ocular correction.

### EEG Recording & Preprocessing

The EEG data were recorded from 65 active electrodes (ActiCap, Brain Products, Munich, Germany) with a sampling rate of 500 Hz using Brain Vision Recorder (Brain Products, Munich, Germany). The electrode Cz was used as the reference during the recording. Four electrodes were placed at the outer canti (LO1, LO2) and below both eyes (IO1, IO2) to record eye movements. Impedances were kept below 5 kΩ. Participants were instructed to avoid movements and blinking during stimulus presentation to minimize muscular and ocular artifacts.

Offline data were re-referenced to average reference and corrected for ocular artifacts using brain electric source analyses (BESA; Ille, Berg, & Scherg, 2002) based on individual eye movements recorded after the experiment. The ocular-corrected EEG data were band pass filtered between 0.5 Hz and 40 Hz and segmented into epochs of 1 s from 200 ms pre-stimulus onset to 800 ms post-stimulus onset. Baselines were corrected to the 200 ms pre-stimulus interval for all segmentations. Segments containing artifacts, hence values ±150 µV or gradients larger than 50 µV were invalidated and thereby excluded from further analysis. EEG data preprocessing was performed in Matlab (The MathWorks Inc.) using customized scripts (Frömer, Maier, & Abdel Rahman, 2018) and EEGLab functions (Delorme & Makeig, 2004).

### Behavioral Data Analysis

Behavioral data were analyzed using linear mixed models (LMMs). LMMs allow for single-trial based parametric analyses and provide estimates of fixed effects, as well as random variance in intercepts and slopes across participants. Analyses were performed in R (R Core Team, 2018) using the lme package (Bates, Maechler, Bolker, & Walker, 2014a). For both *Accuracy* and *RT*, stepwise model reduction was carried out on the respective full model that included all predictors until the reduced model resulted in a significantly decreased fit indicated by the likelihood ratio test and based on the Akaike Information Criterion (AIC) and Bayesian Information Criterion (BIC), which decrease with increasing model fit. Full models included main effects for Scene Congruity, Target Quality and Group, as well as Group by Scene Congruity, Group by Target Quality, and Group by Scene Congruity by Target Quality interactions. The random effects structures comprised participant random intercepts and random slopes for the within-subjects factors Scene Congruity and Target Quality when applicable. To avoid over-parameterization, singular value decomposition was used to determine the random effects structure by eliminating random slopes that prevented model convergence or explained zero variance (Matuschek, Kliegl, Vasishth, Baayen & Bates, 2017).

The final model of *Accuracy* included a binomial generalized linear mixed model (GLMMs) with main effects for Scene Congruity, Target Quality, and Group, as well as Scene Congruity by Group interaction as fixed effects. By-participant random intercepts and random slopes for Scene Congruity and Target Quality were modeled as random effects. *RTs* that were greater or less than 3 standard deviations from the mean of all trials for each subject were considered outliers and excluded from further analyses (4.70% of all trials). The final model of *RT* included main effects of Scene Congruity, Target Quality, and Group, and interactions of Scene Congruity by Group as well as Target Quality by Group. By-participant random intercepts and random slopes for the factor Scene Congruity were modeled as random effects. For all final models, we report estimated effect sizes of the fixed effects, standard errors, z/t-values, and estimates of the square root of the variance components.

### ERP Analysis

Single-trial ERPs for N1, P200 and N400 components were averaged within time windows and electrodes of interest (see Results section) and submitted to LMMs. As with the models for behavior, full models included main effects for Scene Congruity, Target Quality and Group, as well as Group by Scene Congruity, Group by Target Quality and Group by Scene Congruity by Target Quality interactions. For all models, fixed effects structures were determined by model selection using likelihood ratio tests and model fits indicated by AIC and BIC. The random effects structure included random intercepts for participants and random slopes for within-subjects factors, Scene Congruity and Target Quality unless any of those explained zero variance, as determined by singular value decomposition. For all final models, we report estimated effect sizes of the fixed effects, standard errors, t-values, and estimates of the square root of the variance components (Table 2 & 3).

**Table 2.** Statistics of the final model for target detection accuracy rates and for response times.

| **Variable** | **Accuracy** |  |  |  | **RT** |  |  |  |
| --- | --- | --- | --- | --- | --- | --- | --- | --- |
|  | **b** | **SE** | **z** | **p** | **b** | **SE** | **z** | **p** |
| Mean amplitude<br>(Intercept) | 2.51 | 0.07 | 33.55 | <.001 | 0.96 | 0.03 | 36.06 | <.001 |
| Scene Congruity<br>(Incongruent - Congruent) | -0.53 | 0.12 | -4.57 | <.001 | 0.05 | 0.01 | 5.46 | <.001 |
| Target Quality<br>(Blurred - Intact) | -0.49 | 0.12 | -4.07 | <.001 | 0.002 | 0.008 | 0.29 | 0.774 |
| Group<br>(Elderly - Young) | -0.27 | 0.15 | -1.82 | 0.068 | 0.27 | 0.05 | 5.04 | <.001 |
| Congruity: Group | -0.47 | 0.21 | -2.22 | 0.027 | 0.09 | 0.02 | 4.42 | <.001 |
| Quality: Group |  |  | / |  | 0.03 | 0.02 | 1.99 | 0.046 |
| **Variance Component** | **SD** |  |  |  | **SD** |  |  |  |
| Participant | 0.38 |  |  |  | 0.2 |  |  |  |
| Scene Congruity | 0.28 |  |  |  | 0.04 |  |  |  |
| Target Quality | 0.39 |  |  |  | / |  |  |  |
| Residual | / |  |  |  | 0.3 |  |  |  |
| **Goodness of fit** | **Value** |  |  |  | **Value** |  |  |  |
| Log likelihood | -1559.1 |  |  |  | -1115 |  |  |  |
| REML deviance | 3118.2 |  |  |  | 2230 |  |  |  |

Note: Effects and/or variance components not included in the final model are indicated by "/".

**Table 3.** Statistics of the final model for each of ERP components.

| Variable | N1 |  |  |  | P200 |  |  |  | N400 |  |  |  |
| --- | --- | --- | --- | --- | --- | --- | --- | --- | --- | --- | --- | --- |
|  | b | SE | z | p | b | SE | z | p | b | SE | z | p |
| Mean amplitude (Intercept) | 3.98 | 0.5 | 8 | <.001 | 6.92 | 0.46 | 14.98 | <.001 | -1.71 | 0.22 | -7.73 | <.001 |
| Scene Congruity (Incongruent - Congruent) | 0.3 | 0.15 | 1.99 | 0.048 | 0.52 | 0.14 | 3.78 | <.001 | -0.61 | 0.1 | -6.42 | <.001 |
| Target Quality (Blurred - Intact) | 0.63 | 0.17 | 3.75 | <.001 | 0.21 | 0.16 | 1.25 | 0.215 | 0.28 | 0.09 | 3.03 | 0.003 |
| Group (Elderly - Young) | -5.28 | 0.99 | -5.33 | <.001 | -2.59 | 0.92 | -2.81 | 0.006 | 0.48 | 0.44 | 1.09 | 0.28 |
| Congruity: Group | / |  |  |  | -0.54 | 0.28 | -1.96 | 0.049 | / |  |  |  |
| Quality: Group | / |  |  |  | 1.43 | 0.33 | 4.33 | <.001 | / |  |  |  |
| **Variance Component** | **SD** |  |  |  | **SD** |  |  |  | **SD** |  |  |  |
| Participant | 3.81 |  |  |  | 3.54 |  |  |  | 1.68 |  |  |  |
| Scene Congruity | 0.3 |  |  |  | / |  |  |  | 0.21 |  |  |  |
| Target Quality | 0.68 |  |  |  | 0.7 |  |  |  | 0.17 |  |  |  |
| Residual | 5.02 |  |  |  | 4.81 |  |  |  | 3.18 |  |  |  |
| **Goodness of fit** | **Value** |  |  |  | **Value** |  |  |  | **Value** |  |  |  |
| Log likelihood | -14905 |  |  |  | -14690.9 |  |  |  | -12657.6 |  |  |  |
| REML deviance | 29810 |  |  |  | 29381.8 |  |  |  | 25315.3 |  |  |  |

Note: Effects and/or variance components not included in the final model are indicated by "/".

## RESULTS

### Behavioral Results

#### Accuracy

Based on previous studies, we expected that elderly participants would be more sensitive to scene congruity manipulations. In line with our expectation, we observed a significant Group by Scene Congruity interaction [b = -.47, z = -2.22, *p = .027*] on accuracy. The elderly group performed more accurately at detecting the target in congruent than incongruent trials [b = -.79, z = -5.47, *p < .001*], while the young group performed similarly well in both conditions [b = -.26 z = -1.60, *p = .110*], as revealed by a separate follow-up model with Scene Congruity and Target Quality nested within Group (Figure 2b). To verify that the elderly group showed a significantly stronger effect of Scene Congruity, we extracted the random slope estimate of Scene Congruity for each participant and compared the random slopes between the young group and elderly group using a *t-test*. Indeed, the *t-test* revealed a significant difference between the mean random slope for the young group and the elderly group [t = 2.74, *p = .008*], confirming that scene congruity had a greater impact on target recognition in the elderly group relative to the young group. In contrast, both groups were significantly and similarly affected by Target Quality [b = -.49, z = -4.07, *p <.001*] with higher accuracy for intact than blurred images. The mean target detection accuracy of each condition for each group is shown in Figure 2a. While accuracy was numerically slightly lower for the elderly compared to the young group, we found no significant overall difference in accuracy between groups [*b* = -.27, z =-1.82, *p* = .068] when accounting for the Scene Congruency interaction. Taken together, these results indicate that the performance accuracy of the elderly group is more sensitive to Scene Congruity than the young group, but that both groups were similarly sensitive to Target Quality.

**Figure 2.**
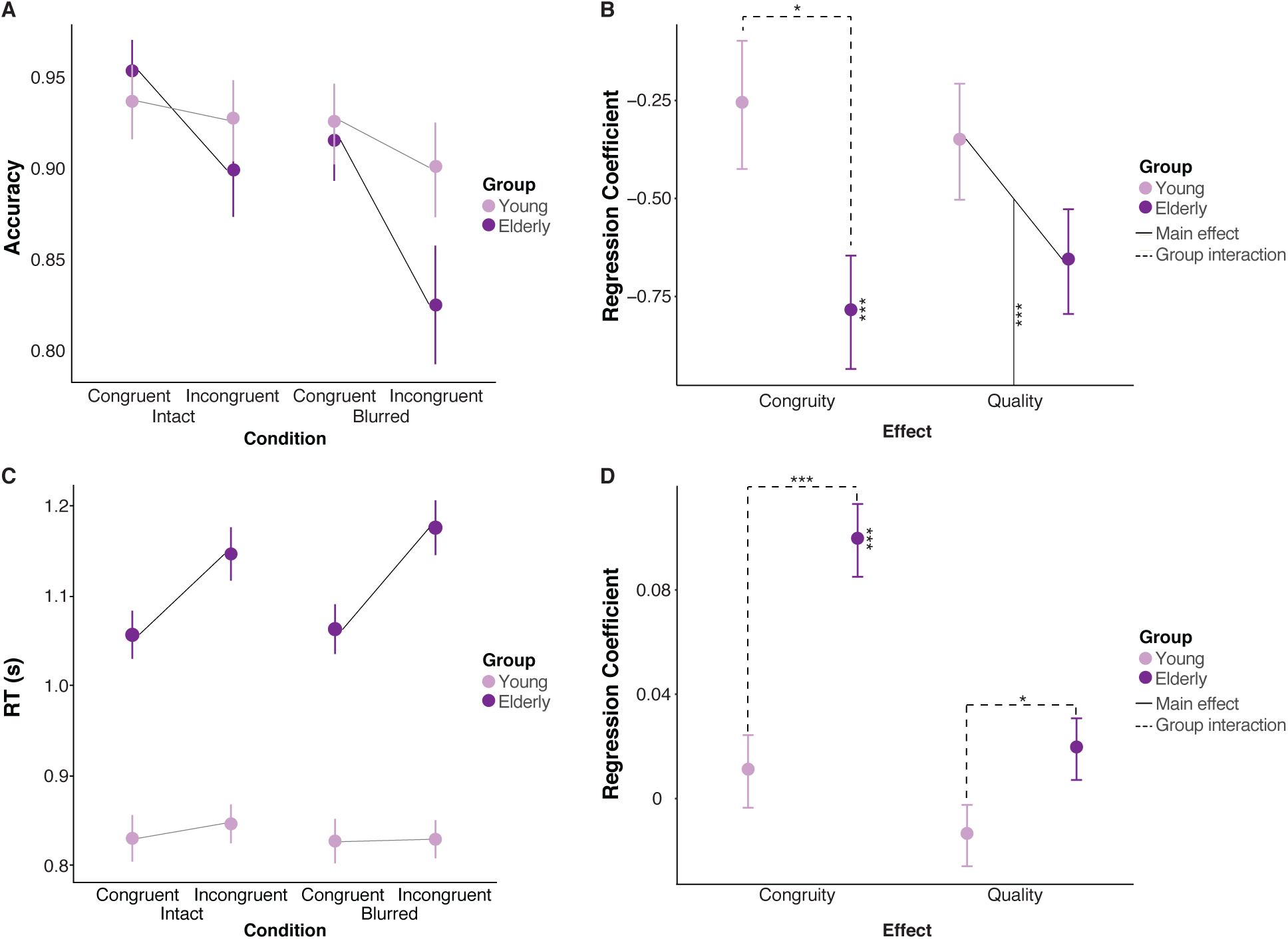
Mean target detection accuracy rates under different conditions for each group (a). Linear mixed-effects model estimates of different effects on accuracy for each group (b). Mean target detection response times under different conditions for each group (c). Linear mixed-effects model estimates of different effects on response time for each group (d). Note: Error bars indicate the respective 95% confidence interval.

#### Reaction Time

We expected that the increased sensitivity to top-down manipulations would be similarly evident in RTs. Indeed, as with accuracy, we observed a significant Group by Scene Congruity interaction [b = .09, t = 4.42, *p < .001*] on RT. RTs were significantly faster on congruent compared to incongruent trials in the elderly group [b = .10, t = 7.10, *p < .001*] but not in the young group [b = .01, t = .75, *p =.458*], as revealed by a separate follow-up model with Scene Congruity and Target Quality nested within Group (Figure 2d). Furthermore, a follow up *t-test* comparing the random slope estimate of Scene Congruity for each participant between the young group and elderly group confirmed that the mean random slope estimate for Scene Congruity was significantly larger in the elderly group compared with the young group, indicating that the impact of Scene Congruity on target recognition increases with age [t = -5.97, *p < .001*]. While a significant Target Quality by Group interaction was also observed [b = .03, t = 1.99, *p = .047*], the effect of Target Quality was not significant within either young group [b = -.01, t = -1.21, p = .228] or elderly group [b = .02, t = 1.60, p = .109] when nesting Target Quality in group. Consistent with previous work (Salthouse, 1996), elderly participants showed overall slower responses compared to young participants [b = .27, t = 5.04, *p < .001*]. The mean RT of each condition for each group is shown in Figure 2c. Taken together, the *RT* results corroborate the *Accuracy* results, and they jointly indicate an age-related increase in the influence of scene congruity on behavioral measures of object recognition.

### ERP Results

The following analyses aimed to shed light on age differences in the time course of bottom-up and top-down processes mediating object recognition. To that end we tested Scene Congruity and Target Quality effects on distinct ERP components involved in different stages of recognition related visual processing. We probed age differences in early sensory and attentional processing during visual discrimination (within 250 ms following stimulus onset) as indicated by the N1 component, subsequent pre-perceptual top-down selective attention processes as indexed by the P200 component, and post-perceptual semantic integration processes as indexed by the N400 component. We expected to see age-related modulation of both N1 and P200 components indicating alterations at the level of perceptual analysis and suppression related top-down control (Gazzaley et al., 2008), respectively. We further predicted interactions between Group and Scene Congruity or Target Quality in these early components due to age-related changes in the dynamics of bottom-up and top-down processing during object recognition. On the other hand, as our hypothesis mainly concerned the changes in the underlying perceptual processes that allow for object recognition in aging, we did not expect to observe age difference in N400, which reflects post-perceptual processing based on semantic knowledge (Ganis & Kutas, 2003; Mudrik et al., 2010).

### N1

The analyses of the N1 component were performed on a predefined posterior ROI (PO9, PO7, O1, O2, PO8 and PO10). Visual inspection of the group grand mean waveforms (Figure 3a) indicated that the N1 peak was delayed in the elderly group relative to the young group. To test whether N1 was indeed significantly delayed, we carried out a peak latency detection procedure in which we first computed the peak N1 latency of each condition for each subject and then analyzed the detected peak latencies in an LMM with the predictors Scene Congruity, Target Quality and Group. This model revealed a significant main effect of Group [b = 36.16, t = 9.15, *p < .001*] revealing that N1 peak latency was delayed by 36 ms in the elderly group compared to the young group [Young: 150.50 ms vs. Elderly: 186.38 ms]. To accommodate the observed age-related delay in subsequent amplitude analyses, the N1 component was quantified separately for the young and elderly group based on the age-adjusted time windows (Young: 150 ms - 180 ms; Elderly: 170 ms - 200 ms). After controlling for the age-related delay in N1 amplitude, we found that elderly participants showed significantly larger N1 amplitudes compared to young participants [b = -5.28, t = -5.33, *p < .001*] (Figure 3c-d). There were no significant interactions between Group and any of the conditions, but across groups we found significantly larger N1 amplitudes both for congruent compared to incongruent scenes [b = .30, t = 1.99, *p = .049*] and for intact compared to blurred targets [b = .63, t = 3.75, *p < 001*] (Figure 4a). In summary, while young and elderly groups displayed similar effects of scene congruity and target quality on N1 amplitudes that is consistent with increased early perceptual analysis for salient bottom-up signals, the elderly group additionally displayed a delayed and larger N1 that is consistent with the need for increased attentional feedback to facilitate the processing of degraded bottom-up signals.

**Figure 3.**
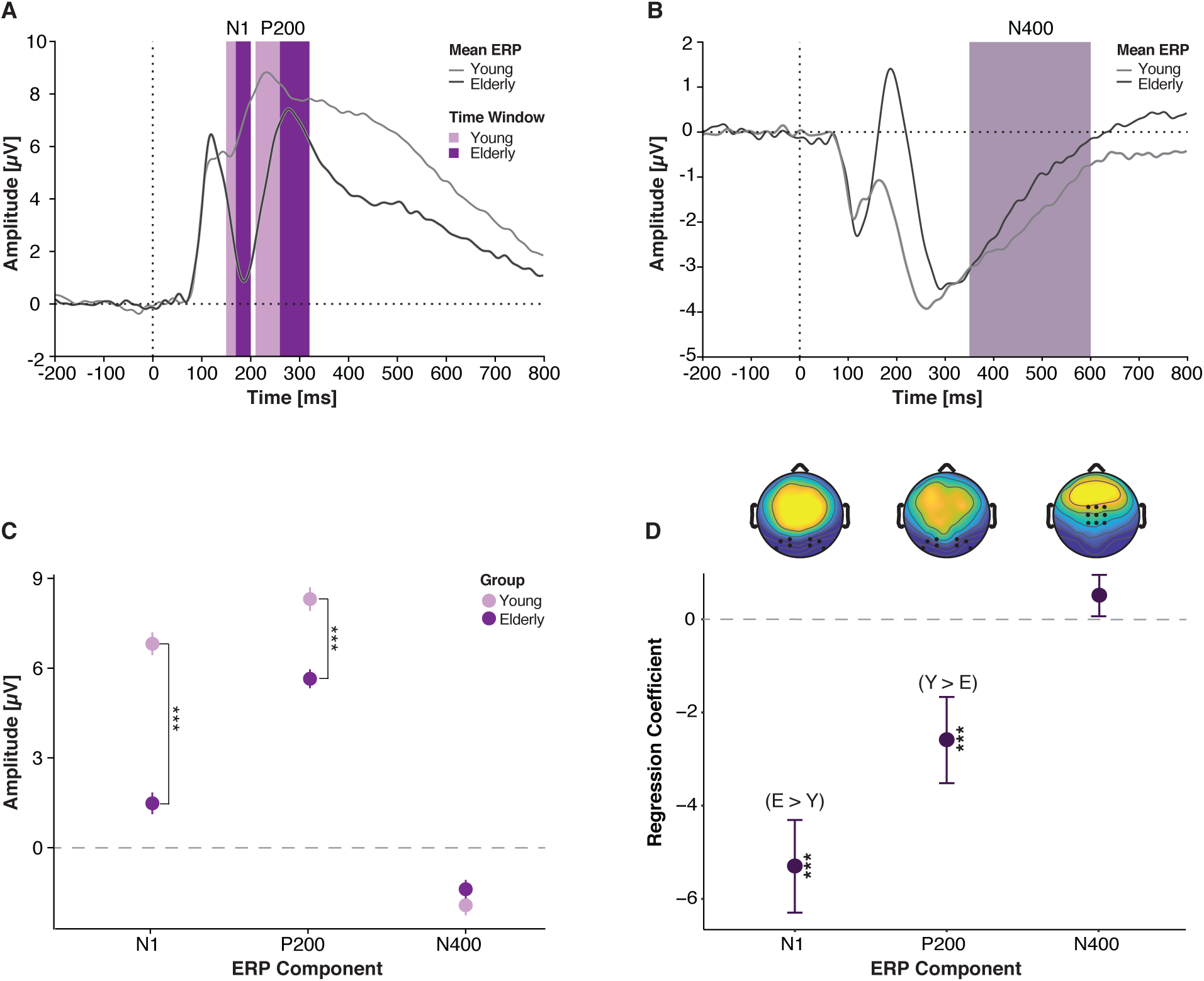
Grand mean amplitude averaged across electrodes within the respective region of interest for N1 and P200 (a), and N400 (b) for each group. Grand mean amplitudes averaged across electrodes within the respective regions of interest for each group (c). Linear mixed effects model estimates of group effect for each ERP component (d). Note (c) (d): Error bars indicate 95% confidence interval. The relative amplitude size between the young group (Y) and the elderly group (E) is denoted for all significant effects.

**Figure 4.**
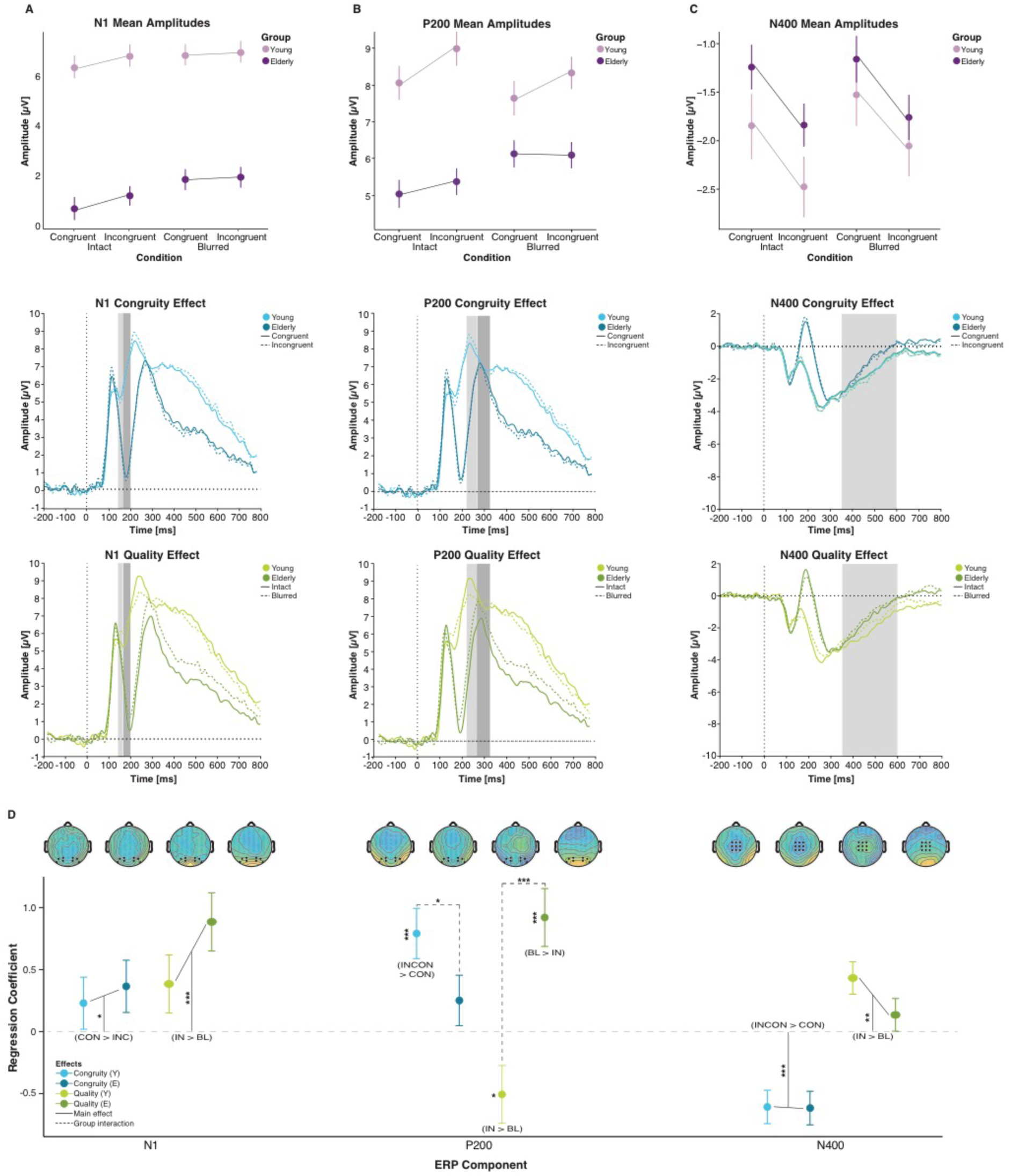
The mean amplitude of each condition for each group (top row), the overall ERP waveform of congruent vs. incongruent condition (second row), and the overall ERP waveform of intact vs. blurred condition (third tow) for N1 (a), P200 (b), and N400 (c). Linear mixed-effects model estimates of congruity and quality effects of each ERP component for each group (d). Note: Error bars indicate 95% confidence interval. The relative amplitude size between congruent (CON) and incongruent (INCON) conditions, and between intact (IN) and blurred (BL) conditions is denoted for all significant effects in (d).

#### P200

The P200 component was quantified at a predefined posterior ROI (PO9, PO7, PO3, O1, O2, PO4, PO8 and PO10). As with the N1, visual inspection of the group grand mean waveforms (Figure 3a) indicated that the P200 peak latency was delayed in the elderly group relative to the young group, and we therefore performed the same peak latency detection procedure and LMM analysis on peak latencies as described above (see N1 in results section). This model revealed a significant main effect of Group [b = 41.15, t = 6.47, *p < .001*] such that P200 peak was delayed by 42 ms in elderly compared to young participants [Young: 250.83 ms vs. Elderly: 292.98 ms]. To examine whether the observed P200 delay might be attributed to the earlier delay in N1, we tested the same LMM on the N1-corrected-latencies of P200 (P200 peak latency - N1 peak latency). Accounting for the N1 delay, there was no significant group difference in P200 latencies [b = 6.20, t = .91, *p = .369*], indicating that the P200 delay resulted from a carryover from the N1 delay. Subsequent amplitude analyses were performed on P200 quantified in the age-adjusted time windows (Young: 210 ms - 270 ms; Elderly: 260 ms - 320 ms).

In contrast to the N1, P200 amplitude was larger in the young group compared to the elderly group [b = -2.59, t = -2.81, *p < .001*] (Figure 3c-d). The groups also significantly differed in their effects of Scene Congruity [b = -.54, t = -1.96, *p = .049*]. A follow up model with the predictors nested within Group revealed a significantly larger P200 amplitudes for incongruent compared to congruent scenes in the young group [b = .79, t = 3.91, *p < .001*] but not in the elderly group [b = .25, t = 1.23, *p = .222*] (Figure 4b). To test whether effects were significantly different in magnitude, we followed up with a *t-test* comparing the random slope estimate of Scene Congruity for each participant between the young group and elderly group, which confirmed that the mean random slope estimate for Scene Congruity was significantly higher in the young compared to the elderly group [t = -4.35, *p < .001*]. Unexpectedly, the groups also differed in the effect of Target Quality [b = 1.43, t = 4.33, *p < .001*]. The nested model revealed that while both groups showed a significant main effect of Target Quality, P200 amplitudes were significantly *reduced* for blurred compared to intact trials in the young group [b = -.51, t = -2.18, *p = .034*], but were significantly *increased* for blurred compared to intact trials elderly group [b = .92, t = 3.88, p < .001] (Figure 4b). We further extracted the random slope estimate of Target Quality for each participant and compared the estimates between the young group and elderly group using a *t-test*, which verified that the two groups differed significantly in the mean random slope estimate for Target Quality [t = -4.35, *p < .001*]. Taken together, the qualitatively distinct patterns of P200 findings displayed by the young and elderly groups are consistent with a reallocation of top-down resources to process unresolved perceptual ambiguity rather than contextual conflict in the elderly, and provide a candidate locus of the observed stronger behavioral susceptibility to scene congruity in elderly compared to young adults.

#### N400

As an index of semantic integration, the N400 component was quantified in a predefined central ROI (FC1, FCz, FC2, C1, Cz, C2, CP1, CPz, and CP2), and in the time window between 350 ms and 600 ms (Figure 3b). Consistent with the N400 effect of semantic violation previously found in language processing (Holcomb, 1993; Kutas & Hillyard, 1980) and scene perception (Barrett & Rugg, 1990; Ganis & Kutas, 2003; Kutas & Federmeier, 2011), both young and elderly group showed a larger N400 amplitude in response to incongruent scenes relative to congruent scenes [b = .61, t = -6.42, *p < .001*] (Figure 4c). Additionally, larger N400 amplitudes were found for intact targets relative to blurred targets [b = .28, t = 3.03, *p = .003*]. However, none of these effects, nor the mean N400 amplitude (Figure 3c-d), differed between groups. These findings suggest that post-perceptual semantic integration is preserved in elderly adults.

### Summary of ERP Results

An overall summary of the LMM estimates of the effects of group, scene congruity and target quality on each ERP component is provided in Figure 4d, and the statistics of the final models for each component are reported in Table 3. Taken together, our findings show that while the efficiency of bottom-up visual processes (as indexed by the N1 component) was altered in elderly compared to young adults, our experimental manipulations affected these processes similarly in both groups. In contrast, top-down regulation of attention (as indexed by the P200 component) was differentially affected by both bottom-up signal integrity and top-down information congruity between the two groups, suggesting a reallocation of top-down resources to process perceptual ambiguity rather than contextual conflict in the elderly group. These findings suggest that observed increased behavioral sensitivity to mismatching contextual information is driven by deficient suppression of irrelevant contextual information. Finally we found that post-perceptual semantic integration at the level of the N400 component was preserved in the elderly group and comparable to the young group.

## DISCUSSION

The present study examined the effects of aging on the neural dynamics between bottom-up and top-down visual processing during a rapid object recognition task by manipulating the strength of both bottom-up (stimulus ambiguity) and top-down (context congruity) signals of the input images. Consistent with previous studies, we found that the behavioral performance of elderly adults (reflected in both accuracy and RT) was more strongly impacted by the presence of incongruent contextual information than young adults. In order to identify the locus of this effect, we examined the impact of these experimental manipulations on the following ERP components indexing object recognition processes at different points along the visual stream: 1) The N1 as an index of early sensory and attentional processing during visual discrimination; 2) The P200 as an index of pre-perceptual top-down selective attention processes; and 3) The N400 as an index of post-perceptual semantic integration processes. As expected, young adults displayed scene congruity effects on all three ERP components, with congruent scenes eliciting larger amplitudes on the N1 and incongruent scenes eliciting larger amplitudes on the P200 and N400. Interestingly, perceptual ambiguity effects were also observed on all three components in young adults, with intact objects consistently eliciting larger amplitudes than blurred objects. More importantly, we observed distinct effects of aging at different stages of visual processing and affecting both bottom-up and top-down recognition processes; these effects arose early at the level of sensory processing and continued to the level of pre-perceptual top-down selective attention, while post-perceptual semantic integration remained intact. Taken together, these finding are consistent with the presence of age-related changes in early interactions between attentional and perceptual processing.

Behaviorally, older adults displayed greater effects of scene congruity on accuracy and RT than did young adults, despite comparable levels of overall recognition accuracy in the two groups. This increased sensitivity to scene congruity may reflect an increased bias in older adults toward additional top-down processing in order to solve object recognition due to age-related declines in bottom-up processing; in this view, the additional top-down engagement may reflect increased effort to leverage contextual information in order to resolve ambiguity within the bottom-up input (Trapp & Bar, 2015). For example, age-related changes in visual processing, including early sensory and perceptual loss (Owsley, 2011; Habak & Faubert, 2000), increased internal noise and decreased signal-to-noise ratio within primary visual cortex (Arena et al., 2013) and neural dedifferentiation within ventral visual stream (Park et al., 2004, 2011; Chee et al., 2006), may lead to increased competition among perceptual hypotheses regarding the object identity during recognition processing that can be resolved through contextual information provided in congruent scenes. Alternatively, the greater recognition impairment displayed by older adults in the presence of mismatching contextual information within incongruent scenes may instead reflect diminished top-down suppression of irrelevant information in older adults (Gazzaley et al., 2008; Zanto et al., 2010). An examination of the age-related changes in the ERP components associated with object recognition may help to clarify the changes in neural dynamics of bottom-up and top-down processing that underpin these behavioral changes.

### Electrophysiological Evidence of Age-Related Changes in Recognition Processing

The electrophysiological findings from the present study indicate that differences between young and elderly groups emerge as early as 150 ms into the course of visual object recognition processing. For both groups, N1 amplitudes were larger for congruent than incongruent scenes and for intact than blurred targets, suggesting that contextually relevant and perceptually intact stimuli provide stronger bottom-up signals that lead to increased early perceptual analysis involving sensory-driven attentional processes as indexed by the N1 (Heinze et al., 1994; Heinze, Luck, Magun, & Hillyard, 1990; Alain & Woods, 1999; Chao & Knight, 1997; Fabiani, Low, Wee, Sable, & Gratton, 2006). The overall amplitude of the N1 was also found to be significantly larger in the elderly group compared to the young group, suggesting that older adults may chronically recruit greater attentional resources during the early stage of visual processing (Heinze et al., 1994; Alain & Woods, 1999; Chao & Knight, 1997; Fabiani et al.,2006) compared to young adults. In addition, an age-related increase in the latency of the N1 may reflect a reduced efficiency of these early attentional mechanisms in older adults (Gazzaley et al., 2008; Zanto et al., 2010). Taken together, these results suggest that while both young and elderly groups demonstrate modulations of the N1 associated with stimulus quality (i.e. target clarity and scene congruity), older adults also preferentially recruit early attentional mechanisms more than young adults in order to compensate for age-related declines in bottom-up visual processing.

In contrast to the quantitative differences (i.e., age-related increases in latency and overall amplitude) observed between young and elderly groups for the N1 component, qualitatively distinct patterns of modulation were observed between the two age groups for the P200 component. Consistent with the role of P200 in facilitating the pre-perceptual processing of relevant information via top-down suppression of irrelevant information (Schapkin, Gusev, & Kuhl, 2000; Potts & Tucker, 2001), young adults displayed larger P200 amplitudes for incongruent than congruent scenes. Young adults also displayed larger P200 amplitudes for intact than blurred targets, suggesting that the demands for suppression of irrelevant information is greater for intact than blurred targets due to the increased saliency of distracting features. Interestingly, older adults did not display the effect of scene congruency on P200 amplitude that was observed in young adults, and they actually displayed the opposite pattern of greater P200 amplitude for blurred than intact targets consistent with an age-related reallocation of selective attention resources to resolve bottom-up ambiguity. We also found a reduction in overall P200 amplitude in older adults, suggesting a decline in executive attention processes with age.

Taken together, these findings suggest that unlike young adults, older adults are not utilizing pre-perceptual top-down selective attention processes to suppress irrelevant information (particularly present in incongruent compared to congruent scenes), but are instead reallocating these attentional resources to address persisting bottom-up perceptual ambiguity issues that were not resolved in early processing stages. That is, because young adults have effectively resolved perceptual ambiguity at earlier processing stages, they can the utilize top-down suppression mechanisms indexed by the P200 to resolve contextual incongruity. In contrast, the inability of the aging visual system to completely resolve perceptual ambiguity in earlier processing stages requires elderly adults to utilize these executive attentional resources to continue resolving object ambiguity rather than scene congruity, suggesting that interference from irrelevant contextual information continues beyond the level of the P200 for the elderly group. Consistent with this view, young adults showed a congruity effect in the P200 but not in accuracy or RT (suggesting that contextual incongruity was resolved at the level of the P200), whereas older adults instead showed congruity effects in both accuracy and RT but not in the P200 (suggesting that contextual incongruity was not resolved at the level of the P200).

Finally, we replicated previous findings of scene congruity effects on N400 amplitude, with larger frontal-central negativity in incongruent than congruent scenes (Demiral, Malcolm, & Henderson, 2012; Ganis & Kutas, 2003; Mudrik et al., 2010; Vo & Wolfe, 2013; Guillaume et al., 2016), in both young and older adults, suggesting that knowledge-dependent processing post-identification (Ganis & Kutas, 2003; Mudrik et al., 2010; Vo & Wolfe, 2013) is preserved with aging. Interestingly, and in contrast to some previous findings of age-related reduction in N400 effect (see Friedmann, 2012 for a review), we did not find age difference in the magnitude of N400 effect. This discrepancy may be due at least in part to differences in the experimental paradigms used to assess the effects of aging on the N400 semantic congruity effect, with previous studies focusing on language processing (e.g. Hamberger & Friedman, 1992; Kutas & Iragui, 1998; Cameli & Phillips, 2000; Federmeier, Van Petten, Schwarz, & Kutas, 2003) rather than on visual scene processing as in the present study. Based on the present results as well as on previous findings indicating that selective top-down suppression impairments displayed by older adults at early stages of visual processing (i.e. within the first 200 ms) recover in later processing stages (Gazzaley et al., 2008), we suggest that in the context of visual recognition, semantic processing post-identification is intact in older adults given that interference from irrelevant information is also reduced later in the time course of visual processing.

Previous electrophysiological studies in older adults have suggested the presence of age-related deficits in top-down suppression within the visual processing stream. Older adults have been found to exhibit disrupted connectivity in frontal-temporal and frontal-occipital regions (Bollinger, Rubens, Zanto, & Gazzaley, 2011; Kalkstein, Checksfield, Bollinger, & Gazzaley, 2011), and this disrupted connectivity has been associated with age-related decline in top-down inhibitory control within visual areas (Gazzaley et al., 2008; Zanto, Toy, & Gazzaley, 2010). On the other hand, a large body of behavioral literature has shown that semantic knowledge is maintained or even increases with age (Abrams & Farrell, 2011; Salthouse, 1993), and that experience-dependent top-down cortical signals may be strengthened with age (Gilbert & Moran, 2016; Kaufman, Keith, & Perlstein, 2006). Top-down feedback activity projected from the OFC has been associated with contextual associations to bottom-up signals, which in turn projects rapid top-down feedback to the posterior regions to improve bottom-up predictions and resolve competition (see Aminoff, Kveraga, & Bar, 2013 for a review). Age-related increase in top-down cortical signals during visual processing (Gilbert & Moran, 2016) may therefore suggest a bias towards context-related top-down processing in older adults. Consistent with this interpretation, increased contextual influence on recognition in older adults has been attributed to extensive visual experiences accumulated throughout lifespan that promotes efficient top-down predictive codes to guide recognition (Aminoff, Gronau, & Bar, 2006; Bar & Aminoff, 2003; Remy et al., 2013; Gilbert & Moran, 2016). The present results, however, did not provide direct support for the notion that context-related top-down facilitation of object recognition improves with age, but rather that context-related facilitation is delayed in older adults in order to further facilitate unresolved perceptual ambiguity. Our failure to find context-related facilitation with aging may be partly due to the random presentation of congruent and incongruent scenes in our experimental design, which has been shown discourage processing strategies relying on contextual information compared to block designs (Whiting, Madden, Pierce & Allen, 2005; Whiting, Madden & Babcock, 2007), a hypothesis that warrants future investigation.

Distinct aging effects at different stages of processing together shift the dynamic interactions between bottom-up and top-down processes mediating object recognition. The present results provide evidence for three important age-related changes in these processes. First, older adults are more dependent on top-down influences (e.g. object-scene associations) for successful object recognition compared with young adults. Second, there is an increase in recruitment of sensory-driven attentional processes in the early stages of visual processing as indexed by N1 in the aging brain, possibly reflecting relatively inefficient bottom-up processing in older adults. Third, older adults show greater reliance on pre-perceptual top-down selective attention processes as indexed by the P200 to address persisting perceptual ambiguity that was not resolved in early processing stages, rather than to resolve higher-order contextual ambiguity. This reallocation of resources (possibly in conjunction with an age-related reduction in executive attention function) leads to an increased sensitivity to interference from irrelevant information that in turn makes object processing particularly vulnerable to incongruent scenes in older adults. Despite these age-related changes in perceptual processing, however, our results also indicate that knowledge-dependent top-down processes concerned with post-perceptual semantic integration is nonetheless preserved in aging.

## Supporting information

Supplemental Model Selection

## REFERENCES

Abdel Rahman, R. A., & Sommer, W. (2008). Seeing what we know and understand: How knowledge shapes perception. Psychonomic bulletin & review, 15(6), 1055–1063.

Abrams, L., & Farrell, M. T. (2011). Language processing in normal aging. The handbook of psycholinguistic and cognitive processes: Perspectives in communication disorders, 49–73.

Alain, C., & Woods, D. L. (1999). Age-related changes in processing auditory stimuli during visual attention: evidence for deficits in inhibitory control and sensory memory. Psychology and aging, 14(3), 507.

Aminoff, E. M., Kveraga, K., & Bar, M. (2013). The role of the parahippocampal cortex in cognition. Trends in cognitive sciences, 17(8), 379–390.

Aminoff, E., Gronau, N., & Bar, M. (2006). The parahippocampal cortex mediates spatial and nonspatial associations. Cerebral Cortex, 17(7), 1493–1503.

Arena, A., Hutchinson, C. V., Shimozaki, S. S., & Long, M. D. (2013). Visual discrimination in noise: behavioural correlates of age-related cortical decline. Behavioural brain research, 243, 102–108.

Auckland, M. E., Cave, K. R., & Donnelly, N. (2007). Nontarget objects can influence perceptual processes during object recognition. Psychonomic bulletin & review, 14(2), 332–337.

Bar, M. (2004). Visual objects in context. Nature Reviews Neuroscience, 5(8), 617.

Bar, M., & Aminoff, E. (2003). Cortical analysis of visual context. Neuron, 38(2), 347–358.

Bar, M., Kassam, K. S., Ghuman, A. S., Boshyan, J., Schmid, A. M., Dale, A. M., … & Halgren, E. (2006). Top-down facilitation of visual recognition. Proceedings of the national academy of sciences, 103(2), 449–454.

Barrett, S. E., & Rugg, M. D. (1990). Event-related potentials and the semantic matching of pictures. Brain and cognition, 14(2), 201–212.

Bates, D., Maechler, M., Bolker, B., & Walker, S. (2014). lme4: Linear mixed-effects models using Eigen and S4. R package version, 1(7), 1–23.

Bollinger, J., Rubens, M. T., Zanto, T. P., & Gazzaley, A. (2010). Expectation-driven changes in cortical functional connectivity influence working memory and long-term memory performance. Journal of Neuroscience, 30(43), 14399–14410.

Bourisly, A. K., & Shuaib, A. (2018). Neurophysiological effects of aging: A P200 ERP study. Translational neuroscience, 9(1), 61–66.

Brainard, D. H. (1997) The Psychophysics Toolbox, Spatial Vision 10:433–436.

Braver, T. S., Barch, D. M., Keys, B. A., Carter, C. S., Cohen, J. D., Kaye, J. A., … & Jagust, W. J. (2001). Context processing in older adults: evidence for a theory relating cognitive control to neurobiology in healthy aging. Journal of Experimental Psychology: General, 130(4), 746.

Chee, M. W., Goh, J. O., Venkatraman, V., Tan, J. C., Gutchess, A., Sutton, B., … & Park, D. (2006). Age-related changes in object processing and contextual binding revealed using fMR adaptation. Journal of cognitive neuroscience, 18(4), 495–507.

Coco, M. I., Nuthmann, A., & Dimigen, O. (2018, March 19). Fixation-related brain activity during semantic integration of object-scene information. Retrieved from osf.io/e78km

Crouzet, S. M., Joubert, O. R., Thorpe, S. J., & Fabre-Thorpe, M. (2009). The bear before the forest, but the city before the cars: revealing early object/background processing. Journal of Vision, 9(8), 954–954.

Dambacher, M., Rolfs, M., Göllner, K., Kliegl, R., & Jacobs, A. M. (2009). Event-related potentials reveal rapid verification of predicted visual input. PLoS One, 4(3), e5047.

Davenport, J. L. (2007). Consistency effects between objects in scenes. Memory & Cognition, 35(3), 393–401.

Davenport, J. L., & Potter, M. C. (2004). Scene consistency in object and background perception. Psychological Science, 15(8), 559–564.

Delorme, A., & Makeig, S. (2004). EEGLAB: an open source toolbox for analysis of single-trial EEG dynamics including independent component analysis. Journal of neuroscience methods, 134(1), 9–21.

Demiral, S. B., Malcolm, G. L., & Henderson, J. M. (2012). ERP correlates of spatially incongruent object identification during scene viewing: Contextual expectancy versus simultaneous processing. Neuropsychologia, 50(7), 1271–1285.

Doran, M. M., & Hoffman, J. E. (2010). The role of visual attention in multiple object tracking: Evidence from ERPs. Attention, Perception, & Psychophysics, 72(1), 33–52.

Drew, T., McCollough, A. W., Horowitz, T. S., & Vogel, E. K. (2009). Attentional enhancement during multiple-object tracking. Psychonomic Bulletin & Review, 16(2), 411–417.

Dyck, M., & Brodeur, M. B. (2015). ERP evidence for the influence of scene context on the recognition of ambiguous and unambiguous objects. Neuropsychologia, 72, 43–51.

Eimer, M. (2000). Effects of face inversion on the structural encoding and recognition of faces: Evidence from event-related brain potentials. Cognitive Brain Research, 10(1-2), 145–158.

Fabiani, M., Low, K. A., Wee, E., Sable, J. J., & Gratton, G. (2006). Reduced suppression or labile memory? Mechanisms of inefficient filtering of irrelevant information in older adults. Journal of cognitive neuroscience, 18(4), 637–650.

Federmeier, K. D., Van Petten, C., Schwartz, T. J., & Kutas, M. (2003). Sounds, words, sentences: age-related changes across levels of language processing. Psychology and aging, 18(4), 858.

Fenske, M. J., Aminoff, E., Gronau, N., & Bar, M. (2006). Top-down facilitation of visual object recognition: object-based and context-based contributions. Progress in brain research, 155, 3–21.

Fize, D., Cauchoix, M., & Fabre-Thorpe, M. (2011). Humans and monkeys share visual representations. Proceedings of the National Academy of Sciences, 108(18), 7635–7640.

Friedman, D. (2012). The components of aging. Oxford handbook of event-related potential components, 1.

Frömer, R., Maier, M., & Abdel Rahman, R. (2018). Group-level EEG-processing pipeline for flexible single trial-based analyses including linear mixed models. Frontiers in Neuroscience, 12, 48.

Ganis, G., & Kutas, M. (2003). An electrophysiological study of scene effects on object identification. Cognitive Brain Research, 16(2), 123–144.

Gazzaley, A., Clapp, W., Kelley, J., McEvoy, K., Knight, R. T., & D’Esposito, M. (2008). Age-related top-down suppression deficit in the early stages of cortical visual memory processing. Proceedings of the National Academy of Sciences, 105(35), 13122–13126.

Gilbert, J. R., & Moran, R. J. (2016). Inputs to prefrontal cortex support visual recognition in the aging brain. Scientific reports, 6, 31943.

Greene, M. R., Botros, A. P., Beck, D. M., & Fei-Fei, L. (2015). What you see is what you expect: rapid scene understanding benefits from prior experience. Attention, Perception, & Psychophysics, 77(4), 1239–1251.

Guillaume, F., Tinard, S., Baier, S., & Dufau, S. (2018). An ERP investigation of object-scene incongruity: The early meeting of knowledge and perception. Journal of psychophysiology, 32(1), 20.

Ille, N., Berg, P., & Scherg, M. (2002). Artifact correction of the ongoing EEG using spatial filters based on artifact and brain signal topographies. Journal of clinical neurophysiology, 19(2), 113–124.

Habak, C., & Faubert, J. (2000). Larger effect of aging on the perception of higher-order stimuli. Vision research, 40(8), 943–950.

Hamberger, M., & Friedman, D. (1992). Event-related potential correlates of repetition priming and stimulus classification in young, middle-aged, and older adults. Journal of Gerontology, 47(6), P395–P405.

Heinze, H. J., Luck, S. J., Mangun, G. R., & Hillyard, S. A. (1990). Visual event-related potentials index focused attention within bilateral stimulus arrays. I. Evidence for early selection. Electroencephalography and clinical neurophysiology, 75(6), 511–527.

Heinze, H. J., Mangun, G. R., Burchert, W., Hinrichs, H., Scholz, M., Münte, T. F., … & Gazzaniga, M. S. (1994). Combined spatial and temporal imaging of brain activity during visual selective attention in humans. Nature, 372(6506), 543–546.

Holcomb, P. J. (1993). Semantic priming and stimulus degradation: Implications for the role of the N400 in language processing. Psychophysiology, 30(1), 47–61.

Joubert, O. R., Fize, D., Rousselet, G. A., & Fabre-Thorpe, M. (2008). Early interference of context congruence on object processing in rapid visual categorization of natural scenes. Journal of Vision, 8(13), 11–11.

Joubert, O. R., Rousselet, G. A., Fize, D., & Fabre-Thorpe, M. (2007). Processing scene context: Fast categorization and object interference. Vision research, 47(26), 3286–3297.

Kalkstein, J., Checksfield, K., Bollinger, J., & Gazzaley, A. (2011). Diminished top-down control underlies a visual imagery deficit in normal aging. Journal of Neuroscience, 31(44), 15768–15774.

Kaufman, D. A., Keith, C. M., & Perlstein, W. M. (2016). Orbitofrontal cortex and the early processing of visual novelty in healthy aging. Frontiers in aging neuroscience, 8, 101.

Kleiner M, Brainard D, Pelli D, 2007, “What’s new in Psychtoolbox-3?” Perception 36 ECVP Abstract Supplement.

Kutas, M., & Federmeier, K. D. (2011). Thirty years and counting: finding meaning in the N400 component of the event-related brain potential (ERP). Annual review of psychology, 62, 621–647.

Kutas, M., & Hillyard, S. A. (1980). Event-related brainpotentials to semantically inappropriate andsurprisingly large words. Biological Psychology, 1980a, 11, 99–116.

Kutas, M., & Iragui, V. (1998). The N400 in a semantic categorization task across 6 decades. Electroencephalography and Clinical Neurophysiology/Evoked Potentials Section, 108(5), 456–471.

Kveraga, K., Boshyan, J., & Bar, M. (2007). Magnocellular projections as the trigger of top-down facilitation in recognition. Journal of Neuroscience, 27(48), 13232–13240.

Lenoble, Q., Bordaberry, P., Rougier, M. B., Boucart, M., & Delord, S. (2013). Influence of visual deficits on object categorization in normal aging. Experimental aging research, 39(2), 145–161.

MATLAB and Statistics Toolbox Release 2017b, The MathWorks, Inc., Natick, Massachusetts, United States.

Matuschek, H., Kliegl, R., Vasishth, S., Baayen, H., & Bates, D. (2017). Balancing Type I error and power in linear mixed models. Journal of Memory and Language, 94, 305–315. doi:10.1016/j.jml.2017.01.001

Mudrik, L., Lamy, D., & Deouell, L. Y. (2010). ERP evidence for context congruity effects during simultaneous object–scene processing. Neuropsychologia, 48(2), 507–517.

Mudrik, L., Shalgi, S., Lamy, D., & Deouell, L. Y. (2014). Synchronous contextual irregularities affect early scene processing: Replication and extension. Neuropsychologia, 56, 447–458.

Oliva, A., & Torralba, A. (2007). The role of context in object recognition. Trends in cognitive sciences, 11(12), 520–527.

Owsley, C. (2011). Aging and vision. Vision research, 51(13), 1610–1622.

Park, D. C., Polk, T. A., Park, R., Minear, M., Savage, A., & Smith, M. R. (2004). Aging reduces neural specialization in ventral visual cortex. Proceedings of the National Academy of Sciences of the United States of America, 101(35), 13091–13095.

Park, J., Carp, J., Kennedy, K. M., Rodrigue, K. M., Bischof, G. N., Huang, C. M., … & Park, D. C. (2012). Neural broadening or neural attenuation? Investigating age-related dedifferentiation in the face network in a large lifespan sample. Journal of Neuroscience, 32(6), 2154–2158.

Payer, D., Marshuetz, C., Sutton, B., Hebrank, A., Welsh, R. C., & Park, D. C. (2006). Decreased neural specialization in old adults on a working memory task. Neuroreport, 17(5), 487–491.

Pelli, D. G. (1997) The VideoToolbox software for visual psychophysics: Transforming numbers into movies, Spatial Vision 10:437-442.

Potts, G. F., & Tucker, D. M. (2001). Frontal evaluation and posterior representation in target detection. Cognitive Brain Research, 11(1), 147–156.

Potts, G. F., Patel, S. H., & Azzam, P. N. (2004). Impact of instructed relevance on the visual ERP. International Journal of Psychophysiology, 52(2), 197–209.

Randolph, C., Tierney, M. C., Mohr, E., & Chase, T. N. (1998). The Repeatable Battery for the Assessment of Neuropsychological Status (RBANS): preliminary clinical validity. Journal of clinical and experimental neuropsychology, 20(3), 310–319.

Rao, R. P., & Ballard, D. H. (1999). Predictive coding in the visual cortex: a functional interpretation of some extra-classical receptive-field effects. Nature neuroscience, 2(1), 79.

Rémy, F., Saint-Aubert, L., Bacon-Macé, N., Vayssière, N., Barbeau, E., & Fabre-Thorpe, M. (2013). Object recognition in congruent and incongruent natural scenes: A life-span study. Vision research, 91, 36–44.

Rossion, B., Gauthier, I., Tarr, M. J., Despland, P., Bruyer, R., Linotte, S., & Crommelinck, M. (2000). The N170 occipito-temporal component is delayed and enhanced to inverted faces but not to inverted objects: an electrophysiological account of face-specific processes in the human brain. Neuroreport, 11(1), 69–72.

Salthouse, T. A. (1993). Speed and knowledge as determinants of adult age differences in verbal tasks. Journal of Gerontology, 48(1), P29–P36.

Salthouse, T. A. (1996). The processing-speed theory of adult age differences in cognition. Psychological review, 103(3), 403.

Schapkin, S. A., Gusev, A. N., & Kuhl, J. (2000). Categorization of unilaterally presented emotional words: an ERP analysis. Acta neurobiologiae experimentalis, 60(1), 17–28.

Serre, T., Oliva, A., & Poggio, T. (2007). A feedforward architecture accounts for rapid categorization. Proceedings of the national academy of sciences, 104(15), 6424–6429.

Störmer, V. S., Li, S. C., Heekeren, H. R., & Lindenberger, U. (2013). Normal aging delays and compromises early multifocal visual attention during object tracking. Journal of cognitive neuroscience, 25(2), 188–202.

Sun, H. M., Simon-Dack, S. L., Gordon, R. D., & Teder, W. A. (2011). Contextual influences on rapid object categorization in natural scenes. Brain research, 1398, 40–54.

Tanaka, J. W., & Curran, T. (2001). A neural basis for expert object recognition. Psychological science, 12(1), 43–47.

Team, R. C. (2018). R: A Language and Environment for Statistical Computing, R Foundation for Statistical Computing, Austria, 2015.

Trapp, S., & Bar, M. (2015). Prediction, context, and competition in visual recognition. Annals of the New York Academy of Sciences, 1339(1), 190–198.

Võ, M. L. H., & Wolfe, J. M. (2013). Differential electrophysiological signatures of semantic and syntactic scene processing. Psychological science, 24(9), 1816–1823.

Voss, M. W., Erickson, K. I., Chaddock, L., Prakash, R. S., Colcombe, S. J., Morris, K. S., … & Kramer, A. F. (2008). Dedifferentiation in the visual cortex: an fMRI investigation of individual differences in older adults. Brain research, 1244, 121–131.

Whiting, W. L., Madden, D. J., & Babcock, K. J. (2007). Overriding age differences in attentional capture with top-down processing. Psychology and Aging, 22(2), 223–232.

Whiting, W. L., Madden, D. J., Pierce, T. W., & Allen, P. A. (2005). Searching from the top down: Ageing and attentional guidance during singleton detection. The Quarterly Journal of Experimental Psychology Section A, 58(1), 72–97.

Zanto, T. P., Toy, B., & Gazzaley, A. (2010). Delays in neural processing during working memory encoding in normal aging. Neuropsychologia, 48(1), 13–25.

