## Supplemental Model Selection for "Age-Related Changes in the Neural Dynamics of Bottom-Up and Top-Down Processing During Visual Object Recognition: An Electrophysiological Investigation"

### Linear Mixed Effects Model Selection Procedure

## N1

N1 amplitude was first analyzed with the full model including Scene Congruity, Target Quality and Group as well as all possible two-way and three-way interactions as predictors. This full model revealed no significant interactions among the predictors. A stepwise reduction of the interaction terms resulted in the final model with Scene Congruity, Target Quality and Group as fixed effect structure and by-participant intercepts along with random slopes for Scene Congruity and Target Quality as random structure. The final model yielded better model fit indices [AIC: Full model = 29841 vs. Final model = 29832; BIC: Full model = 29965 vs. Final model = 29903] and did not fit significantly worse than the full mode as indicated by the likelihood ratio test [ΔΧ^^^2(8) = 6.58, *p = .58*].

## P200

P200 amplitude was first analyzed with the full model including Scene Congruity, Target Quality and Group as well as all possible two-way and three-way interactions as predictors. All non-significant interaction terms were removed using the stepwise exclusion procedure, resulting in the final model with Scene Congruity, Target Quality, Group as well as Group by Scene Congruity and Group by Target Quality interactions as fixed effect structure and by-participant intercepts along with random slopes for Target Quality as random structure. The final model indicated better model fit indices [AIC: Full model = 29416 vs. Final model = 29402; BIC: Full model = 29467 vs. Final model = 29540] and did not fit significantly worse than the full mode as indicated by the likelihood ratio test [ΔΧ^^^2(9) = 3.27, *p = .95*].

## N400

N400 amplitude was first analyzed with the full model including Scene Congruity, Target Quality and Group as well as all possible two-way and three-way interactions as predictors. This full model revealed no significant interactions terms which were subsequently removed in the stepwise reduction procedure as described above. The final model included Scene Congruity, Target Quality and Group as predictors along with by-participant intercepts and random slopes for Scene Congruity and Target Quality as random structure. The final model showed better model fit indices [AIC: Full model = 25337 vs. Final model = 25337; BIC: Full model = 25409 vs. Final model = 25415] and did not fit significantly worse than the full mode as indicated by the likelihood ratio test [ΔΧ^^^2(1) = 2.45, *p = .12*].
